# Top-down, auditory pallial regulation of the social behavior network

**DOI:** 10.1101/2023.03.08.531754

**Authors:** Jeremy A. Spool, Anna P. Lally, Luke Remage-Healey

**Affiliations:** Neuroscience and Behavior, Center for Neuroendocrine Studies, University of Massachusetts, Amherst, MA 01003, USA

**Keywords:** cortico-hypothalamic pathway, aromatase, dual role, competing motivations, immediate early gene, courtship signal

## Abstract

Social encounters rely on sensory cues that carry nuanced information to guide social decision-making. While high-level features of social signals are processed in the telencephalic pallium, nuclei controlling social behaviors, called the social behavior network (SBN), reside mainly in the diencephalon. Although it is well known how mammalian olfactory pallium interfaces with the SBN, there is little information for how pallial processing of other sensory modalities can modulate SBN circuits. This is surprising given the importance of complex vocalizations, for example, for social behavior in many vertebrate taxa such as humans and birds. Using gregarious and highly vocal songbirds, female Zebra finches, we asked to what extent auditory pallial circuits provide consequential input to the SBN as it processes social sensory cues. We transiently inactivated auditory pallium of female Zebra finches during song playback and examined song-induced activation in SBN nuclei. Auditory pallial inactivation impaired responses to song specifically within the lateral ventromedial nucleus of the hypothalamus (VMHl), providing the first evidence in vertebrates of a connection between auditory pallium and the SBN. This same treatment elevated feeding behavior, which also correlated with VMHl activation. This suggests that signals from auditory pallium to VMHl can tune the balance between social attention and feeding drive. A descending influence of sensory pallium on hypothalamic circuits could therefore provide a functional connection for the integration of social stimuli with internal state to influence social decision-making.

**Significance:** Sensory cues such as vocalizations contain important social information. These social signals can be substantially nuanced, containing information about vocalizer identity, prior experience, valence, and emotional state. Processing these features of vocalizations necessitates processing the fast, complex sound streams in song or speech, which depends on circuits in pallial cortex. But whether and how this information is then transferred to social circuits in limbic and hypothalamic regions remains a mystery. Here, we identify a top-down influence of the songbird auditory pallium on one specific node of the social behavior network within the hypothalamus. Descending functional connections such as these may be critical for the wide range of vertebrate species that rely on intricate sensory communication signals to guide social decision-making.

## Introduction

Our sensory environment is a key part of our social lives. For example, in many social animals like humans and birds, dynamic vocal exchanges shape interactions and impact subsequent encounters. In the brain, social cues such as vocalizations are processed hierarchically across several brain regions from hindbrain to the pallium. In other areas of the brain, several limbic and hypothalamic nuclei are critically involved in the control of social behaviors; these ‘social behavior’ nuclei are densely interconnected, enriched in steroid signaling, and are often referred to as the Social Behavior Network (SBN) (1, 2). ‘Social behavior’ nuclei such as those in the SBN are thought to control social behavior in part by integrating external social signals, like vocalizations, with internal state. In order to understand social behavioral control, it is essential to understand how socially-relevant cues, once fully unpacked by sensory systems, reach social behavior circuits to modulate their output.

It has long been known that SBN nuclei respond to all major sensory modalities tested (e.g., olfaction: (3–5); auditory: (6–11); somatosensory: (12–14); visual: (15–17)), suggesting that rich connections with sensory systems are a critical aspect of how these nuclei participate in the control of social behavior (18, 19). Social stimulus selectivity in sensory processing areas (20–26), taken together with the rich sensory representations in the SBN laid out above, appear to blur the lines between regions delineated as strictly sensory processing and those that control social behavior (18). This is most evident in studies that directly link the social behavioral impacts of olfactory processing in the pallium of mammals to specific SBN circuit nodes. (1,27–34). For example, in mice, juvenile pheromones processed in olfactory cortex (i.e., the vomeronasal organ) decreased sexual receptivity in females; this sensory-guided social decision-making relied on SBN nuclei (31). For mammals, network models and schematics of social circuits now regularly include specific olfactory pallial areas (18, 33, 35).

When it comes to other sensory modalities commonly used across vertebrate species, such as vision and audition, there is less known about circuits that transform sensory representations into social behavior. For the auditory system, brain areas relatively ‘early’ in the ascending auditory pathway clearly provide input to the SBN. Comparative work in frogs, birds, and rodents, have revealed influences of the auditory midbrain and thalamus on circuits that control social behavior in the SBN (11, 36–41). However, auditory signals in many species are complex, and are not fully processed until reaching higher order circuits such as the auditory pallium. In many vertebrate taxa, the auditory thalamus sends a large projection to a primary recipient area of auditory pallium (including the cortex in mammals); this primary area then projects to a number of secondary auditory pallial areas (42–46). In mice, inactivation or lesion of primary auditory cortex impacts pup retrieval in mothers but not virgin females (47, 48). In birds, lesion studies of auditory pallium alter female preferences for male vocalizations, disrupt pairbond formations, and can alter entire social networks (49–51). Auditory pallium is clearly instrumental to fully unpack sociosensory stimuli to dissect auditory scenes, extract meaning from complex auditory signals (i.e., language, song), learn new sounds, and recognize conspecifics (25, 52–57). These functions are vital to social interactions, so these higher-order sensory percepts must eventually reach circuits that control social behavior.

To uncover a top-down influence of auditory pallium on circuits controlling social behavior, birds are an excellent study system. Zebra finches (*Taeniopygia guttata*) live in large social groups and can remember upwards of forty individuals by their vocalizations alone (58), and the auditory pallium of this species has been well-characterized (e.g., (21, 43, 49, 52, 59–66). In birds, current evidence suggests that auditory pallium is functionally connected to SBN nuclei. Auditory-evoked immediate early gene expression between secondary auditory pallium and SBN nuclei is highly correlated in female Zebra finches (7), and secondary auditory pallium projects to potential intermediate nuclei, which may then project to the SBN (67, 68).

We asked whether auditory pallium is necessary for auditory responses observed in SBN nuclei, consistent with a top-down auditory pallial influence on circuits that control social behavior. Alternatively, auditory pallium could process sensory cues in parallel to the SBN, and in this way both systems could guide behaviors via downstream effector regions. In this latter case, sensory representations in SBN nuclei would not depend on auditory pallium. We transiently inactivated auditory pallium of female Zebra finches during male song playback, and examined song-induced immediate early gene activation in SBN nuclei. Our findings identify the first known functional connection between auditory pallium and a specific node of the SBN, the lateral ventromedial nucleus of the hypothalamus.

## Results

### Auditory pallial inactivation suppressed activity in a downstream auditory pallial region

To determine whether inactivation of primary auditory pallium disrupted coding in downstream auditory regions, we injected GABA receptor agonists (baclofen & muscimol) centered on primary auditory pallium (i.e., Field L) and recorded extracellularly from the caudomedial nidopallium (NCM), a secondary auditory pallial region downstream of Field L. Prior to injection in Field L, multiunit traces in NCM displayed characteristic baseline and stimulus-response properties as in previous work (e.g., (69); Fig. 1B). In multiunit traces, NCM is characterized by irregular spontaneous activity and heightened activity during the duration of natural acoustic sounds such as bird songs and calls. Upon baclofen & muscimol injection into Field L, after 5 minutes spontaneous NCM activity was entirely altered, qualitatively shifting to intermittent bursts of activity in between periods of inactivity. With respect to stimulus-driven activity, song playback now evoked brief onset responses to some syllables, and overall evoked firing was suppressed. We recorded at this site at three further time points, 10, 17, and 60 minutes, to assess the duration of the effect GABA receptor agonists in Field L had on NCM. A linear mixed model revealed a main effect of time of recording on song-evoked multiunit firing rates (F_(4,93)_ = 14.89, p < 0.001), with responses 5 min before GABA receptor agonist infusion in Field L differing from all other time points (compared to 5 min post infusion, t_92_ = 7.30, p < 0.0001; to 10 min post infusion, t_92_ =5.78, p < 0.0001; to 17 min post infusion, t_92_ = 4.06, p = 0.0003; to 60 min post infusion, t_92_ = 4.05, p = 0.0003; Fig. 1C). Song-evoked multiunit firing rates slowly increased between the 5 min and 60 min recording (5 min post infusion to 60 min post infusion; t_92_ = −3.25, p = 0.0027), but the qualitative change in baseline activity and song-evoked firing rates to stimuli did not fully recover (Fig. 1C). Recordings at various sites in NCM as the electrode was retracted confirmed that this suppression was evident for at least 400 μm dorsal to the recording site. Thus, inactivation of primary auditory pallium using GABA receptor agonists leads to suppression of downstream auditory processing regions for at least 60 min.

**Figure 1.**
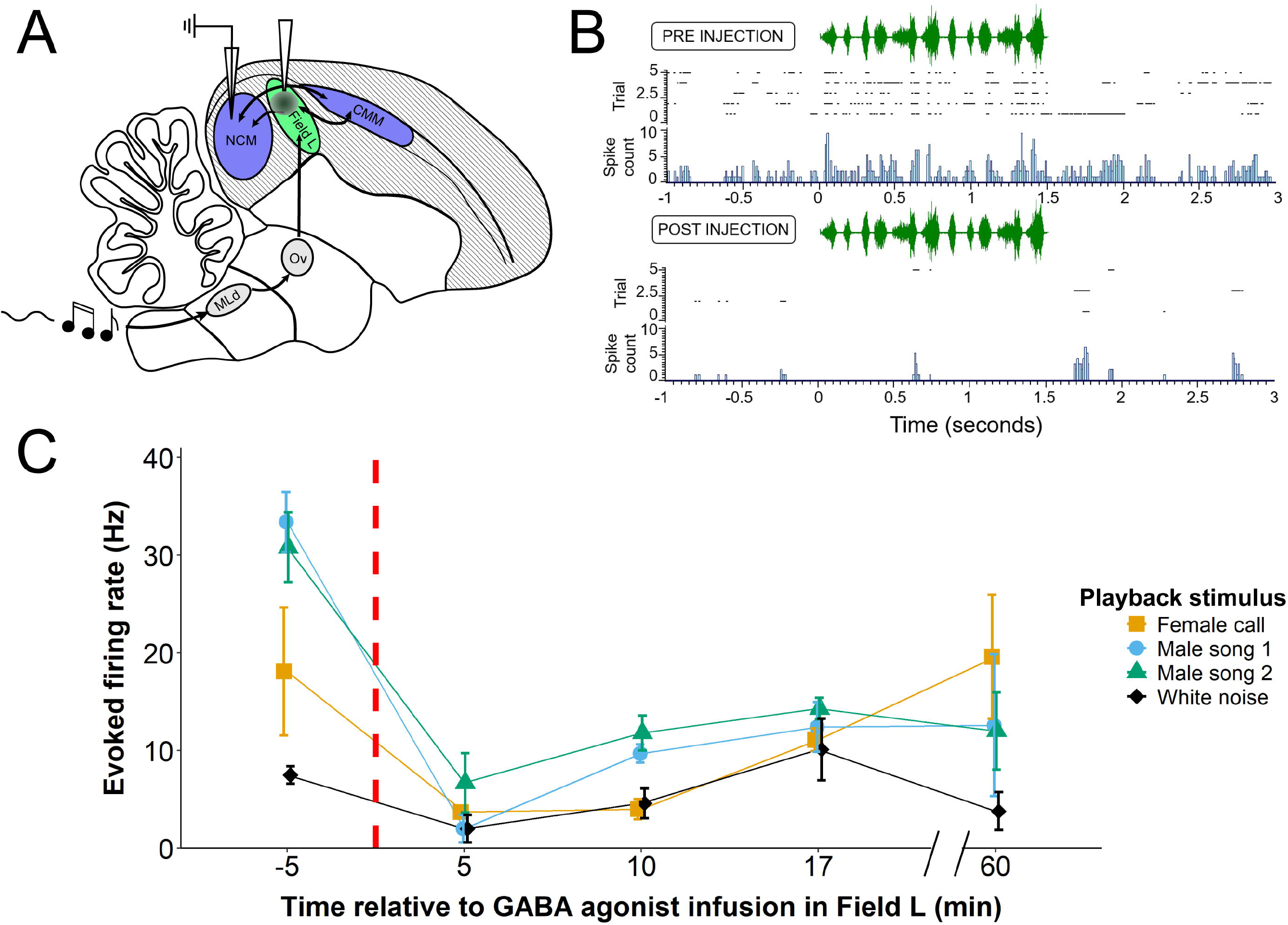
Successful targeting of primary auditory pallium, Field L, with GABA receptor agonists results in impaired auditory processing in a downstream pallial region. (**A**) A sagittal schematic of the Zebra finch brain, and the major route of auditory information flow to the pallium (shaded) from subpallium (gray). Prior to, and following, an infusion of baclofen & muscimol (GABA receptor agonists), in primary auditory pallium (Field L; shown in green), extracellular physiological recordings were made in a downstream secondary auditory pallial region, the caudomedial nidopallium (NCM). Secondary auditory regions (NCM, and caudomedial mesopallium (CMM)) are shown in blue. Subpallial auditory regions (mesencephalicus lateralis, pars dorsalis (MLd) and nucleus ovoidalis (Ov)) are shown in gray). (**B**) Raster plots and summary histograms of the multiunit response in NCM before (top) and five min following (bottom) an infusion of baclofen & muscimol into upstream Field L. A sonogram of the Zebra finch song played beginning at time 0 is depicted in green above each plot. (**C**) Mean multiunit evoked firing rates to various auditory stimuli recorded at various time points before and after baclofen & muscimol (GABA receptor agonists) infusion into Field L. The vertical red dashed line represents the time of infusion.

### GABA receptor agonists in auditory pallium disrupted the immediate early gene response to song specifically in the lateral ventromedial nucleus of the hypothalamus

Next, we asked how disruption of auditory pallium would impact downstream nuclei in the social behavior network. To do this, we targeted bilateral cannulas centered on dorsal primary auditory pallium, Field L. We confirmed cannula placement across treatment groups (see Materials and Methods for detail; Fig. 2A,B). We asked whether song exposure, GABA receptor agonist infusion in auditory pallium, or the interaction of these two treatment levels, affected the number of egr-1 cells labeled in our brain regions of interest in the Social Behavior Network (Fig. 2C-E; Fig. 3A). For the lateral ventromedial nucleus of the hypothalamus (VMHl), a two-way ANOVA revealed a significant main effect of song treatment (F_1,13_ = 5.98, p = 0.029), and a significant interaction between song treatment and cannula infusion treatment (F_1,13_ = 17.68, p = 0.001). Consistent with previous accounts that VMHl is auditory responsive (6), Tukey HSD postdoc tests revealed a significant song playback effect for birds that received a saline infusion in auditory pallium; birds that heard song had greater numbers of egr-1 cells compared to birds that didn’t (p = 0.002; Fig. 3D). By contrast, there was no such difference for birds that received GABA receptor agonist infusion in auditory pallium; birds that heard song had equivalent numbers of egr-1 cells compared to birds that didn’t. Unexpectedly, birds that received GABA receptor agonist infusions in auditory pallium, regardless of song exposure, had egr-1 cell counts somewhere in between birds that received saline infusions. Tukey HSD tests showed that birds treated with silence and GABA receptor agonist fusion in auditory pallium had higher levels of egr-1 in VMHl compared to birds treated with silence and saline infusion (p = 0.008; Fig. 3D). No other groups were statistically different from one another (all other p > 0.1; Fig. 3D).

**Figure 2.**
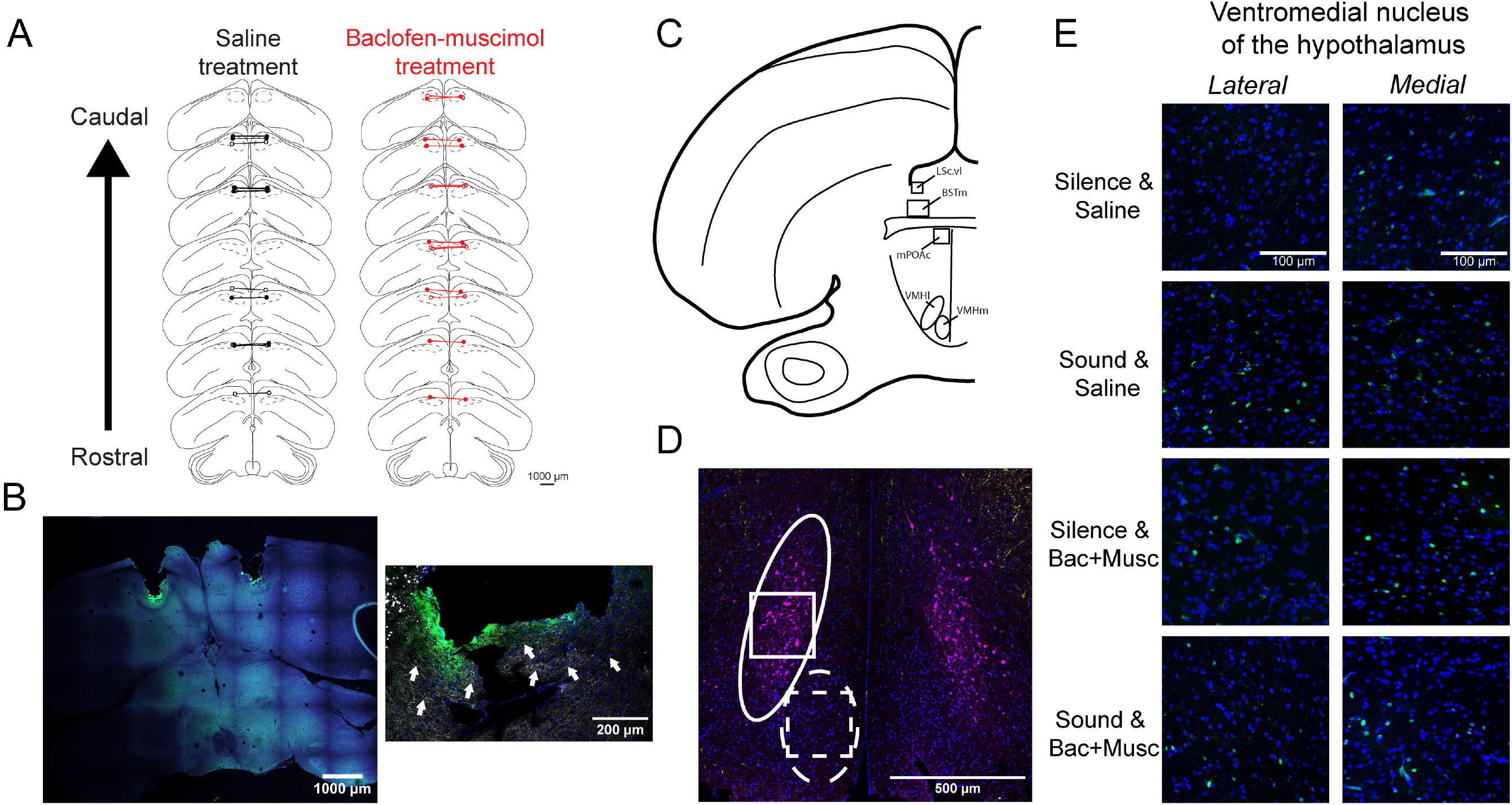
Targeting of cannulas and regions of interest. (**A**) Rostral-to-caudal map of bilateral cannula barrel tip locations at maximum depth in animals receiving saline infusions (left series) or baclofen & muscimol infusions (right series). Playback condition is indicated by open circles (silence condition) versus closed circles (song playback condition). (**B**) (*Left*) Exemplar whole section image of cannula tracks at maximum depth. Green around cannula tip represents Alexa 488 fluorophore in infusion. Blue is used for contrast. (*Right*) A close up of a cannula barrel tip imaged for DAPI (blue), Alexa 488 (green) and myelinated fibers (white; imaged using the reflective properties of myelin, see materials and methods; white arrows indicate examples of fibers). (**C**) Coronal plane depicted regions of interest quantified for egr-1 labeling. LSc.vl = lateral septum, caudoventrolateral; BSTm = bed nucleus of the stria terminalis, medial; mPOAc = medial preoptic area, caudal; VMHl and VMHm = ventromedial nucleus of the hypothalamus, lateral and medial respectively. (**D**) Bilateral view of the ventromedial nucleus of the hypothalamus (VMH) in a coronal plane. The lateral VMH is defined as the extent of the population of cells expressing the enzyme aromatase (solid outline; anti-aromatase = magenta), and the medial VMH is an ovoid nucleus ventromedial to VMHl (dashed outline). DAPI (blue) and parvalbumin (gold) are used as counterstains. (**E**) Exemplar images of egr-1 expression (green) and DAPI (blue) in the VMHl (left column) and VMHm (right column) taken from the positions of boxes in B. Each row represents an experimental treatment group, including whether a bird received a saline infusion or baclofen & muscimol infusion (Bac+Musc; GABA receptor agonists) into auditory pallium, and whether a bird was exposed to silence or song playback.

**Figure 3.**
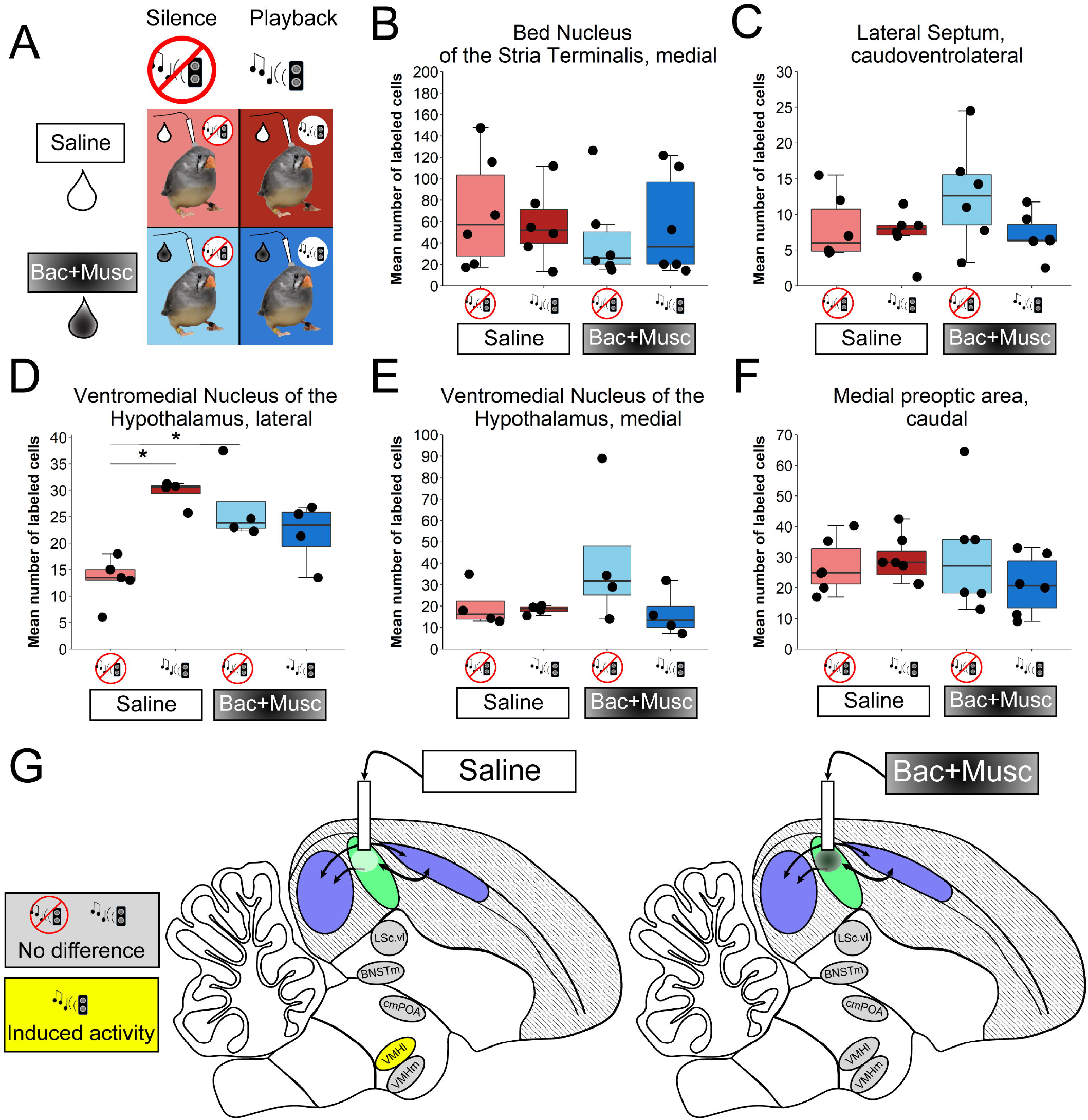
Effects of sound and cannula infusion treatments on egr-1 cell counts. (**A**) Schematic of experimental treatment groups. Female zebra finches received either saline or baclofen & muscimol (Bac+Musc; GABA receptor agonists) in auditory pallium. Then, birds were either exposed to silence or playback of conspecific male song. (**B-F**) Box plots of cell counts of tissue immunolabeled for egr-1 in multiple regions of the Social Behavior Network. * = p< 0.05. (**G**) Green area represents primary auditory pallium, and blue areas represent downstream auditory secondary auditory pallium. (*Left*) When birds receive a saline infusion in auditory pallium, egr-1 expression in the lateral ventromedial nucleus of the hypothalamus (VMHl) is higher in birds exposed to playback than birds exposed to silence. (*Right*) When birds receive a GABA receptor agonist infusion in auditory pallium, there is no difference in egr-1 expression in VMHl between silence and playback-exposed birds.

No other brain regions measured, including neighboring medial VMH (VMHm), had a significant difference in egr-1 cell counts among the 4 treatment groups (including the medial bed nucleus of the stria terminalis (BSTm), the caudoventrolateral lateral septum (LSc.vl), or the caudal medial preoptic area (mPOAc); all p > 0.12 for all main effects and interactions in 2-way ANOVA tests in each region; Fig. 3B,C,E,F)). Thus, these findings reveal a clear descending influence of the auditory pallium on a specific nucleus in the Social Behavior Network, the lateral VMH (Fig 3G).

### GABA receptor agonist infusion in auditory pallium increased foraging behavior

Next, we examined how experimental treatments affected behavior to ascertain whether cannula inactivation of auditory pallium influenced how birds attend to sounds, as well as other homeostatic-related behaviors. We compared four behavioral measures between groups, including attentive behaviors (alert postures and head tilting), beak gaping (a behavior sometimes observed following handling stress), and foraging (i.e., feeding) behavior (Fig. 4A-D). A 2-way ANOVA revealed a significant main effect of cannula infusion treatment in auditory pallium on bouts of foraging behavior (F_1,19_ = 16.5, p = 0.0007; Fig. 4D). Tukey’s HSD posthoc tests showed that birds receiving GABA receptor agonist infusions in auditory pallium engaged in significantly more foraging bouts than birds given saline infusions in auditory pallium (p = 0.0008; Fig. 4D). These postdoc tests revealed no significant main effect of song playback treatment nor an interaction between song and cannula infusion treatments (all p > 0.08; Fig. 4D). No other significant differences in behavior were identified across treatments.

**Figure 4.**
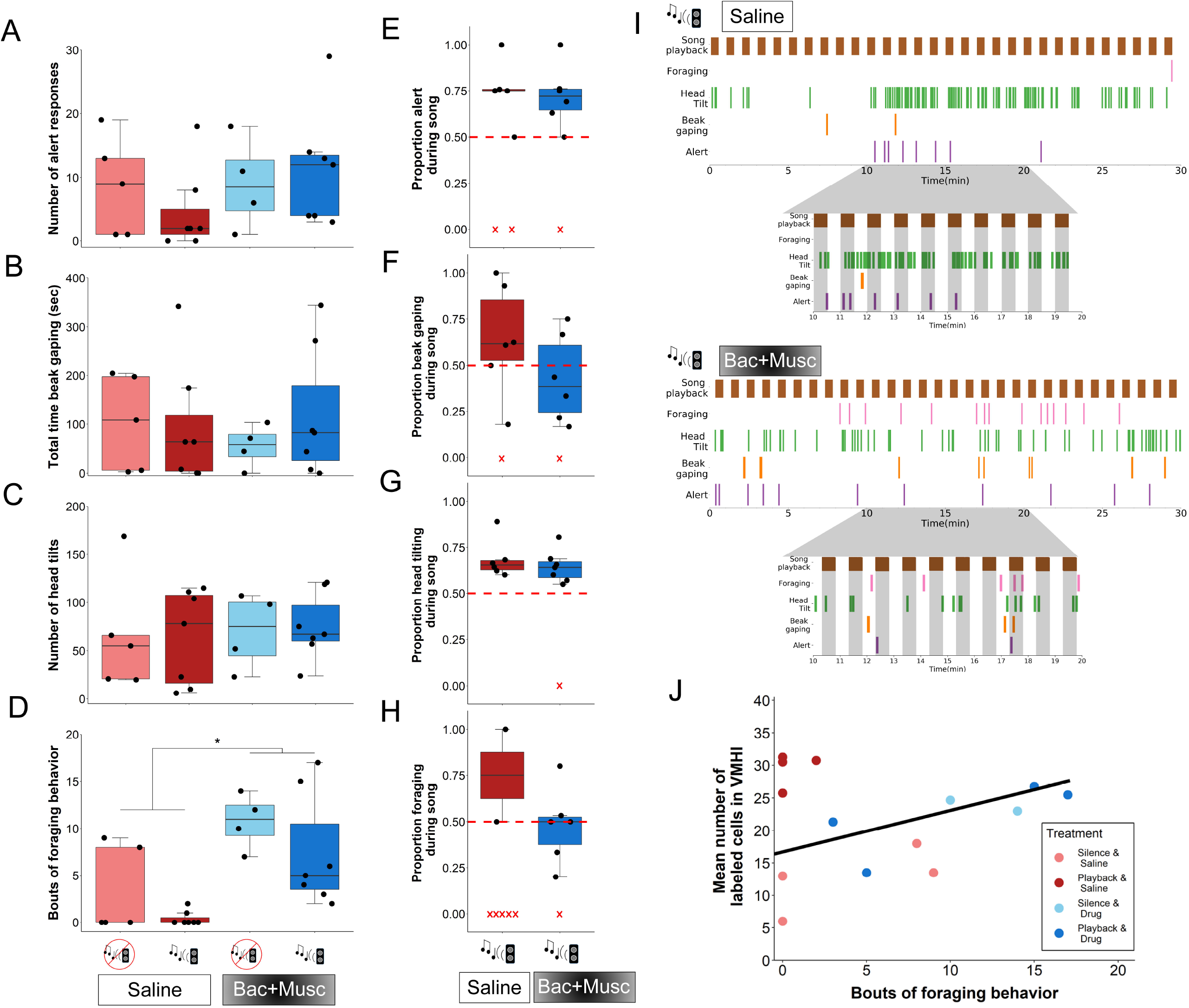
Effects of sound and cannula infusion treatments on number of (**A**) alert responses, (**B**) total time beak gaping, (**C**) number of head tiles, and (**D**) bouts of foraging behavior. Bac+Musc = baclofen & muscimol (GABA receptor agonists). (**E-H**) For birds that heard song playback, we also calculated the proportions of behavior specifically when song playback was on compared to when song playback was off. The red dashed line at 0.5 denotes where 50% of a behavior was during song playback periods. Red ‘X’s at zero on the Y-axis denote animals that did not perform the behavior during the paradigm; because these animals by definition cannot have a proportion of behavior performed during either song or silence, these animals were excluded from the analysis. (**I**) Exemplar ethograms over the 30 min period of song playback for a bird with saline infused in auditory pallium (top) and baclofen & muscimol (Bac+Musc) infused in auditory pallium (bottom). The inset below each ethogram is an enlargement or the ethogram between 10 and 20 minutes. Shaded columns are bounded by the duration of song playback periods to demonstrate alignment of behavior during versus outside song. (**J**) Scatterplot of foraging behavior and egr-1 immunolabeling in the lateral ventromedial nucleus of the hypothalamus (VMHl). Colors represent experimental treatment group. Regression line represents a significant relationship (p = 0.005) between foraging behavior and VMHl egr-1 immunolabeling when controlling for treatment group. * = p< 0.05.

It was unexpected that GABA receptor agonist infusions in auditory pallium affected foraging behavior; we reasoned that disruption of high-level auditory signal processing may have led birds to divert time allocation to increases in other behaviors like foraging. Because our song playback was structured as repeated periods of playback and silence (30 sec playback alternating with 30 sec silence), this allowed for a comparison of whether cannula infusion treatment altered behaviors specifically during the playback periods. We compared the proportions of each behavior’s occurrence during song playback periods versus silent periods. If GABA receptor agonists in auditory pallium affected general attentiveness to playback, then alert behaviors and head tilts would be higher during song playback periods in saline-treated birds as compared to GABA receptor agonist-treated birds during song playback. Instead, there were no differences in proportion of behaviors during song periods compared to silent periods (Fig 4E-H), including alert behaviors, head tilting, beak gaping, or foraging (with the caveat that only two saline-treated birds foraged at all; Welch’s t-tests all p > 0.2). In both groups, birds performed more alert behaviors (Saline infusion and Song: t_4_=3.18, p = 0.034; GABA receptor agonist infusion and Song: t_5_=3.27, p = 0.022) and head tilts (Saline infusion and Song: t_5_=5.27, p = 0.003; GABA receptor agonist infusion and Song: t_6_=5.05, p = 0.002) during song compared to silence (Fig. 4E,G,I). Whereas, the number of foraging behaviors and beak gaping did not differ during song or silence (Fig. 4F,H; all p > 0.3). Neither the number of attentive behaviors, or their timing, then, explain the increase in foraging behavior caused by infusion of GABA receptor agonists in auditory pallium.

### Auditory pallial input disrupts a relationship between VMHl egr-1 expression and foraging behavior

The specificity with which GABA receptor agonists in auditory pallium affected both foraging behavior as well as egr-1 expression in VMHl recalls previous literature that demonstrated the involvement of VMH in feeding behavior in rodents and birds (70–75). We asked whether individual variation in egr-1 induction in VMHl was explained by changes in foraging behavior. We ran a 2-way ANCOVA, with foraging behavior, song treatment, and cannula infusion treatment, as well as the song X cannula infusion treatment interaction, as predictor variables. The addition of foraging behavior as a covariate did not change the effect of treatment on egr-1 expression in VMHl as described above (Fig. 3D; song treatment (F_1,9_ = 28.30, p <0.001), and the interaction between song and cannula infusion treatment (F_1,9_ = 25.95, p <0.001)). However, in addition, foraging behavior significantly predicted egr-1 expression in VMHl (F_1,9_ = 7.40, p = 0.024). Visual inspection of the relationship between foraging behavior and egr-1 cell counts in VMHl by experimental group revealed that all birds that received song playback and saline infusion in auditory pallium were clustered apart from all other birds (Fig. 4J). Posthoc simple linear regressions confirmed this observation (simple linear regression including all birds, not controlling for experimental treatment (F_1,12_ = 0.12, p = 0.73, Adjusted R^2^ = −0.07); simple linear regression excluding the clustered Silence & Saline treated birds (F_1,8_ = 14.41, p = 0.005, Adjusted R^2^ = 0.60; regression line in Fig. 4J). By contrast, a 2-way ANCOVA revealed no effect of experimental treatment or foraging behavior on egr-1 expression in nearby VMHm (all p > 0.17). These findings suggest a functional role for auditory pallial inputs selectively to VMHl. Specifically, when auditory pallial processing is dampened or absent, egr-1 expression in VMHl reflects foraging drive.

## Discussion

Research over the past few decades has demonstrated context-dependent auditory responses in nuclei critical for social behavioral control (6–10). This auditory information is critical for social decision-making, and is processed, in part, by the auditory pallium (25, 52–57). In the present study, we reveal a functional link from auditory pallium to a specific hypothalamic nucleus critical for social behavior, the VMHl. The VMH is implicated in the control of multiple social behaviors across vertebrate species (35, 76–78), but also involved in aspects of homeostasis, including feeding state and fat regulation (71, 73–76, 79), similar to other hypothalamic and preoptic regions that have dual roles in social behavior and homeostatic functions (80, 81). In the present study, inactivation of auditory pallium increased foraging behavior. Furthermore, foraging behavior correlated with immediate early gene expression in the VMHl. Thus, we provide evidence that VMHl (*1*) receives information about social auditory cues processed by the pallium and (*2*) reflects feeding drive when auditory pallial input is dampened or absent. Descending connections from auditory pallium to the hypothalamus may therefore be critical pathways for social decision-making.

At present the available data leave open the possibility for both direct and indirect pathways from auditory pallium to VMHl. In a recent study focused on VMHm (68), retrograde tracing identified sparse inputs from NCM, a secondary auditory pallial region, but it is unclear to what extent VMHl is included in these direct descending projections. There are, at least, three likely possibilities for indirect pathways from auditory pallium to VMHl. One example is through the shell of the auditory thalamus. This shell pathway projects to VMHl (68), though it is unclear whether this shell area receives descending projections from auditory pallium. In mammals, a similar shell structure around the auditory thalamus has emerged as a candidate “secondary thalamic” area (82). Also in mice, a multimodal thalamic nucleus that receives descending input from auditory cortex sends a major projection to the paraventricular nucleus of the hypothalamus (41). Another likely possibility is via amygdalar regions such as the medial arcopallium in birds, which projects to VMH and is auditory-responsive ((6, 68, 83); also see (67) for one possible multisynaptic pathway through the ventral arcopallium and the mesolimbic reward system). Furthermore, the medial acropallium is also thought to be analogous to the mammalian medial amygdala (84–86); in mammals the medial amygdala is thought to route salient olfactory sensory information from pallium to the VMH to guide decision-making (87). Finally, prefrontal cortex may be important in conveying highly processed sensory information to the hypothalamus in general (e.g., (88, 89)), though no major efferent pathways to the hypothalamus from the avian analog have yet been described. An important next step will be using intersectional tracing approaches to identify brain regions that link auditory pallium and the VMHl.

An intriguing aspect of our data is that GABA receptor agonist treatment in auditory pallium, regardless of song treatment, increased egr-1 expression in VMHl. First, this emphasizes that pharmacological inhibition of auditory pallium without any auditory stimulus is sufficient to reveal the functional connection from auditory pallium to VMHl. One possibility is that top-down auditory inputs to VMHl are normally inhibitory. Disrupting auditory pallium may disinhibit VMHl, leading to more general excitability. Another complimentary explanation is that, with GABA receptor agonists in auditory pallium, egr-1 expression in VMHl reflects neuronal activity associated with other behaviors. Both foraging behavior and egr-1 in VMHl were elevated by GABA receptor agonist infusion into auditory pallium. Furthermore, at the individual level, foraging behavior was directly correlated with egr-1 expression in VMHl. Because there is a known role for VMHl in feeding motivation and metabolism (70, 71, 73–76), our findings now indicate that auditory input from pallium shapes VMHl activity in neuronal populations that integrate homeostatic and social drives. We observed no qualitative topographic separation of cells in birds with the highest feeding drive compared to birds with song-induced egr-1, suggesting that the same cells either participate in both social and feeding circuitry depending on the context, or that social and feeding cells are interspersed populations. Data from this present work generate new hypotheses for circuit mechanisms by which hypothalamic nuclei influence motivational state depending on social sensory input.

Our paradigm detected an effect of song on egr-1 expression specifically in the lateral VMH. This is consistent with a previous study in female White-throated sparrows that showed, without estradiol treatment, that VMHl was the only SBN nucleus that differentiated songs from tones (6). Our data do not, however, suggest VMHl is the only subpallial nucleus critical for social behavior that receives input from auditory pallium. Maney et al. 2008 (6) detected several nuclei with auditory responses in the SBN framework that were not detected in the present study. This may be due partly to the choice of study species. Zebra finches are socially-monogamous and opportunistic breeders that form pair-bonds, whereas White-throated sparrows are seasonally-breeding and highly territorial songbirds. Thus, in our paradigm, female finches may have evaluated stranger song for reproductive salience, but this type of stimulus in this species may not have activated cell populations in other SBN regions that may reflect social vigilance, anxiety, aggression, etc. Also, although we focused on regions defined by the SBN framework, there are other regions not included in our study that are critical for social decisionmaking (e.g., see frameworks that expand focus beyond the SBN (18, 90), or proposals to reconsider these network frameworks entirely (91)). Future studies can expand upon the richness of social stimuli and naturalistic contexts and also examine males in these contexts as well, as there may be sex differences in auditory pallial input to nuclei involved in social behavioral control.

Auditory pallium also receives information from the hypothalamus to modulate sensory representations. In mammals, oxytocin neurons in the periventricular nucleus of the hypothalamus project to the primary auditory cortex to modulate auditory responses to infant vocalizations (47). In birds, the medial preoptic area, through indirect projections to the pallial premotor nucleus HVC, modulates the motivation to produce courtship song (92, 93). These observations, in combination with the present study, predict the occurrence of bidirectional loops between higher pallium and hypothalamus and limbic regions in the SBN that continuously modulate sensory processing in concert with motivational state.

In developing the SBN framework, Sarah Newman originally used data from male rodents to propose that social decision-making emerged from the integration of internal and external cues (1). In this context, ‘external cues’ referred to coding of social scents, while ‘internal cues’ referred to sex steroid signaling that altered motivational states within the SBN. In a variety of systems, nuclei within this SBN framework are now known to process a wide array of sensory cues, including interoceptive signals (3–10, 12–17, 81, 94, 95). Thus, both external and internal sensory inputs to brain regions that control social behavior are critical components of social circuitry. Yet in many cases such ‘sensory-social’ brain pathways remain in a black box. In this study in female songbirds we have identified two end-points of such a pathway connecting auditory pallium and the VMHl. How recipient nuclei like VMHl use this auditory information could depend on other coincident external information and internal states represented in either auditory-recipient cells or non-auditory neighboring cell groups, such as feeding drive. These findings highlight the importance of expanding our understanding of social circuits to include ‘upstream’ pallial structures from sensory modalities critical for social interactions.

## Materials and Methods

### Animals

We used female Zebra finches (*Taeniopygia guttata*) in this study housed in unisex aviaries under a photoperiod of 14-h light: 10-h dark. Birds were in acoustic and visual contact with a neighboring unisex aviary of male birds. Food and water were provided ad libitum, and birds received weekly dietary enrichment (e.g., egg food, fresh millet branches, cuttlebone). For electrophysiology, N = 1 female was used to confirm downstream network effects of pharmacological manipulation of Field L. N= 28 females were used in cannulation surgeries and subsequent behavioral testing and immediate early gene labeling. All procedures and protocols adhered to the guidelines of the National Institutes of Health Guide for the Care and Use of Laboratory Animals, and were approved by the University of Massachusetts, Amherst Institutional Animal Care and Use Committee.

### Electrophysiology and pharmacology verification

We performed extracellular recordings on N = 1 bird *in vivo* under urethane anesthesia, following methods in previous studies (69). To administer urethane anesthesia, injections of 30 μL 20% urethane were made in the pectoral muscle every 45 min (specific amount depended on the mass of the bird), totaling 90-120 μL. We then moved the anesthetized bird to a custom stereotaxic apparatus, where we performed a crainiotomy to expose the brain surface dorsal to Field L and the caudomedial nidopallium (NCM; a secondary auditory region downstream of Field L), and fixed a stainless steel headpost to the head using acrylic cement. Immediately following, we moved the bird to a sound-attenuation booth (Industrial Acoustics) on an air table (TMC, Peabody, MA) where the bird was fixed to a custom stereotax (Herb Adams Engineering) at a 45° head angle using the attached headpost. Experiments were conducted in the left hemisphere.

Coordinates used were in reference to the caudal edge of the bifurcation of the sagittal sinus with the head tiled at a 45° angle (Field L = 1.8 rostral, 1.2 lateral, 1.7 ventral; NCM = 1.1 rostral, 0.7 lateral, 1.75 ventral). Prior to recordings, a glass pipette (tip diameter: 30 μm) was loaded with mineral oil and attached to a Nanoject III (Drummond Scientific Company, Broomall, PA). Using the Nanoject, a solution of GABA receptor agonists was drawn into the pipette tip, containing 1 mM baclofen, 0.1 mM muscimol, and 5% fluorescent dextran amines (3,000 MW; Life Technologies Corporation, Carlsbad, CA) in 0.9% saline. The Nanoject was attached to one of two micromanipulators (World Precision Instruments w/ Kantetec, Sarasota, FL), and the glass pipette was lowered to reach Field L. Using the other micromanipulator, a tungsten electrode (A-M Systems, Sequim, WA) was lowered into NCM.

We conducted auditory playback experiments as in previous studies (69, 96), using 3 conspecific vocalizations (1 contact call; directed songs from 2 different unfamiliar males) and white noise, pseudorandomly presented 5 times each, with an interstimulus interval of 10+2 sec. Recordings were made using the tungsten electrode in NCM. After two playback trials, we injected 300 nL of the GABA receptor agonist solution into Field L. 5, 10, 17, and 60 minutes following the injection in Field L, we conducted auditory playback experiments and collected recordings in NCM at the same site. Following the experiment, while retracting the tungsten electrode from the brain, we performed qualitative checks of response of other NCM sites to auditory playback to confirm the observed responses were not an artifact of the chosen site. Following recordings, the bird was transcardially perfused and the brain was extracted for anatomical confirmation of the Field L injection site (localized using fluorescence in injection cocktail).

Recordings were amplified, bandpass filtered (300 to 5000 Hz; A-M Systems), and digitized at 16.67 kHz (Micro 1401, Spike2 software; Cambridge Electronic Design). Data were processed in Spike2 (version 7.04). Recordings were thresholded by the experimenter using a level crossed only by high-amplitude events (i.e., 2-3x the noise band), excluding all small-amplitude events, as in previous work (e.g., (62, 97)). Peristimulus histograms and raster plots were generated to examine multiunit responses to auditory stimuli.

### Cannulation surgeries

One week prior to test day, we implanted N = 28 female zebra finches with bilateral cannulas (Plastics One; 2.4 mm center-to-center distance) targeting Field L, the primary thalamorecipient of ascending auditory signals in the avian pallium. Birds were fasted ~30 min prior to surgery, anesthetized with 2% isoflurane (VetOne) in 2 L/min O2, fixed to a custom stereotax (Herb Adams Engineering) equipped with a heating pad (DC Neurocraft) at a 45° head angle, and maintained on 1.5% isoflurane, 1 L/min O2 for the duration of the surgery. Points overlying Field L were marked by scoring the skull lateral and rostral to our coordinates (i.e., a crosshair), and a craniotomy exposed the brain surface (Field L coordinates = 1.8 mm rostral, 1.2 mm lateral of stereotaxic zero, defined as the caudal edge of the bifurcation of the midsaggital sinus). Dummy bilateral cannula were fitted into guide cannula, and lowered 1.4 mm below the brain surface, just dorsal to the main myelinated fiber tracks terminating in Field L (as shown in Fig. 2B). Metabond (C&B) was applied to the skull around the cannula as an anchor for subsequent application of dental cement, which secured the guide cannula to the skull.

### Testing paradigm

24 hours before the experiment began, birds were habituated with a mock injection, which mimicked the handling and time course of a real injection. Birds were isolated in an acoustic chamber overnight with a playback speaker and a video camera (Sony), and all experiments were run the following day between 8:30 AM and 11 AM. Birds were given one of four treatments in a factorial design: Silence (i.e., heard no song), Saline infusion; Silence, GABA receptor agonist infusion (Baclofen (1 mM) and Muscimol (0.1 mM) as described above) dissolved in saline infusion; Song (i.e., playback of conspecific zebra finch songs), Saline infusion; Song, GABA receptor agonist infusion (N = 7 per treatment; N = 28 birds total). All solutions included 0.2% Alexa 488 fluorophore to aid in visualizing the site of infusion (Life Technologies, Carlsbad, CA). Birds were run in a randomized block design; treatments were randomized among cohorts of 4 birds at a time (but within each cohort, each treatment combination was represented once).

For infusion, the dummy cannula was removed and replaced with an infusion cannula. In all treatments, we infused 300 nL using 15 μL Hamilton syringes over the course of 1 min, driven by a Harvard Apparatus pump (Harvard Apparatus, Hollison, MA), followed by a 2 min wait to allow solution to diffuse. The infusion cannula was then withdrawn and replaced with the dummy cannula. Tubing filled with ddH20 connected the syringes to the infusion cannula. We confirmed flow through the cannula by following the meniscus between the treatment solution and the ddH20 filling the rest of the tubing. The infusion procedure took approximately 5 minutes from the time the bird was caught to when it was returned to the chamber. Song playback or silence commenced 5 min after the bird was returned to the chamber, and lasted 30 min. Following these 30 min, we waited an additional 30 min to allow for egr-1 expression (6, 98). During this time, birds were video recorded for offline quantification of behavior. Birds were then rapidly captured, overdosed on isoflurane, and transcardially perfused with approximately 30 mL of chilled 0.01 M phosphate buffered saline (PBS), followed by approximately 30 mL of chilled 4% paraformaldehyde.

Two birds passed away in between cannula infusion and trial day, leaving sample sizes at: Silence/Saline, N = 6; Song/Saline, N = 7; Silence/Baclofen&Muscimol, N = 6; Song/Baclofen&Muscimol, N = 7.

### Tissue processing and Immunofluorescence

Brains were dissected from the skull and left overnight in 4% paraformaldehyde, then transferred to dehydrate in a 30 % sucrose in 0.01 M PBS until the brain sank. We then froze brains by immersing them in cryo-embedding compound (Ted Pella Inc., Redding, CA) in 2 x 2 x 2 inch plastic blocks at −80 °C. Brains were sectioned in the coronal plane in a cryostat (Leica) at 30 μm. Sections were stored at −20 °C in cryoprotectant solution (30 % sucrose, 30 % ethylene glycol, 1 % polyvinylpyrrolidone, in 0.1 M phosphate buffer).

To verify cannula placement, every fourth section was collected onto glass slides and coverslipped with Prolong antifade mounting medium with a fluorescent stain for DAPI (Invitrogen, catalog #P36962). Cannula tracks were identified under a light microscope (Zeiss) and proximity of the track to Field L was first confirmed through comparison to available Zebra Finch brain atlases (99, 100). We then used a confocal microscope (Nikon) to assess the presence of Alexa fluorophore at the cannula tips. As an additional confirmation of placement, we layered a second image on the same microscope using spectral confocal reflectance (SCoRe) microscopy to visualize the dense myelin that characterizes Field L2a in this region of the pallium (99, 101, 102). In all but 2 cases, cannula barrel tips were either within the boundaries of Field L or just dorsal to it (i.e., within 300 microns dorsal of fibers; see Fig. 2A). In the 2 cases that cannula barrel tips were greater than 300 microns dorsal to Field L fibers, barrel tips were within the ventral caudal mesopallium, a part of auditory pallium. For one of the two animals, which received a saline infusion, analyses were unchanged by its inclusion or removal. For the other, which received a baclofen & muscimol infusion, this animal was identified in outlier analyses in every brain region as having > 2 SD egr-1 expression compared to other animals in its group, and was thus removed independent of consideration of cannula placement (see statistics below).

We conducted fluorescent immunolabeling to detect immediate early gene expression and protein markers that aid in defining brain regions of interest. Tissue sections were washed 5 times for 5 min each in 0.01 M PBS, 3 times for 10 min each in 0.01 M PBS with 0.3% triton (PBT), blocked for 1 hr using 10 % normal donkey serum, and incubated for two days at 4 °C in primary antibodies mixed in blocking serum. Tissue sections were run in two series: one series was incubated with rabbit anti-egr1 at a dilution of 1:1,000 (RRID: AB_2231020) and mouse anti-parvalbumin at a dilution of 1:10,000 (RRID: AB_2174013); the second series was incubated with rabbit anti-aromatase at a dilution of 1:2,000 (aromatase antibody provided as a generous gift from Dr. Colin Saldanha) and mouse anti-tyrosine hydroxylase at a dilution of 1:2,000 (RRID: AB_572268).

Sections were then moved to room temperature and washed 3 times for 10 min each in 0.1 % PBT, followed by a 1 hr incubation in secondary antibodies at a dilution of 1:500 in 0.3 % PBT. Secondary antibodies used included donkey anti-rabbit Alexa 488 (RRID: AB_2556546), and donkey anti-mouse 594 (RRID: AB_2556543). From the secondary antibody step forward, tissue was covered to prevent photobleaching of fluorescence. Sections were finally washed 3 times for 10 min each in 0.1 % PBT, mounted onto subbed microscope slides (Fisher), and coverslipped using Prolong antifade mounting medium with a fluorescent stain for DAPI (Invitrogen, catalog #P36962). Slides dried overnight at room temperature, and were stored at 4 °C until imaging.

### Imaging and Regions of Interest

Sections were imaged on a confocal microscope (A1SP; Nikon, Tokyo, Japan) at the UMass light microscopy core facility. Images were acquired using NIS-Elements software (RRID: SCR_002776). We determined gain and laser intensity separately for each tissue section to minimize background fluorescence.

We focused our analysis on three nodes of the social behavior network that have previously exhibited song-induced egr-1 expression in female songbirds, and 2 regions that did not, based on previous work (6). Regions measured include the ventrolateral subdivison of lateral septum (LSvl), the medial bed nucleus of the stria terminalis (BSTm), the the caudal medial preoptic area (mPOAc; also abbreviated as POM in other bird literature), and the medial and lateral subdivisions of the ventromedial nucleus of the hypothalamus (VMHm and VMHl). For all regions, we took 10 μm Z stacks with 2 μm steps at either 20x or 40x. Brain regions were localized using landmarks established in previous literature ((6, 17, 68); Fig. 2C,D). For LS, the area quantified was a square with 195 μm sides. For BSTm, the area quantified was a rectangle 400 μm by 300 μm. For mPOAc, the area quantified was a square with 318 μm sides. For VMH, we took Z stacks across tiled large images (3 wide by 4 tall) with 5% stitching. To define subregions of VMH, we used guidance from previous literature (6, 68). VMHm is a compact ovoid nucleus at the level of anterior commissure that sits above the optic tract. The area quantified was an ellipse 500 μm on the dorsal/ventral axis and 250 μm on the medial/lateral axis (area = 98 μm^2^). For VMHl, we quantified the area dorsolateral to VMHm, the extent of which was determined based on aromatase fluorescent immunolabeling on alternative sections (Fig. 2D). The area quantified was an ellipse 800 μm on the max axis, and 250 microns on the minimum axis (area = 157 μm^2^); using the max axis at 90 degrees as a reference point, the ellipse was tilted approximately +15 degrees to match the slope of the aromatase positive population on alternate sections.

### Data analysis

#### Electrophysiology and pharmacological verification

Mean multinunit firing rates were calculated in Hz for each auditory stimulus across trials at each time point relative to baclofen & muscimol infusion into Field L.

#### Behavior

Two observers unaware of experimental treatment quantified behavioral videos for the following behaviors: alert responses, which were defined by a quick straightening of posture in which the bird lengthened it’s neck; head tilts, which were defined as a head movement that changed the angle of the head relative to the ground; beak gaping, which was defined as beak opening not associated with foraging (both number of bouts, and bout length, was quantified); foraging, which was defined as pecking at either the food bowl, or directed at objects such as spilled seed on the cage floor. Video camera issues prevented quantification in 3 cases, making sample sizes for behavior as follows: Silence/Saline, N = 5; Song/Saline, N = 7; Silence/Baclofen&Muscimol, N = 4; Song/Baclofen&Muscimol, N = 7. One observer quantified all videos, and the second observer quantified 75%of videos – videos scored by both observers were used to confirm consistency of scoring. Each observer scored each video without audio. Following behavioral scoring, observers recorded timestamps for when individual playback periods began and ended; these timestamps were used later to test whether behavior in experimental treatments that heard playback differed between silent periods and playback periods. Observers quantified behavior for each animal for 35 minutes: this included the 5 min period immediately following Field L infusion, as well as the following 30 min of playback or silence. Behavior from all 35 min was used to assess group differences in the total number of a given behavior. For analyses assessing the proportion of behavior during song playback or silence, we used behavior from the 30 min period of playback to control for the amount of time animals spent listening to song playback VS sitting in silence.

#### Cell counting

Cells positive for egr-1 immunolabeling were quantified using Nikon NIS Elements at the UMass light microscopy core facility. Automatic detection of positive cells in NIS Elements was curated using the following parameters: 1) A threshold above background levels of fluorescence, 2) Volume of label across Z-stacks (defined by range of DAPI nuclei volume in the same brain region), and 3) Sphericity (to filter out elongated objects). To ascertain the quality of automated counts, a person, unaware of treatment condition and of the result of automated counts, manually performed cell counting in a subset of images in each brain region.

Outliers were identified if a data point was greater than 2 standard deviations from the mean of an experimental group. One outlier was identified in this process in all brain regions measured (i.e., the same individual). This, as well as tissue damage during brain processing that made some regions unquantifiable, leads to the final sample sizes for each brain region, which are presented below. For BSTm egr-1, N = 6 per group. For LSc.vl egr-1, N = 6 per group. For cPOM egr-1, N = 7 for the Sound/Saline group, and N = 6 for all other groups. For VMHm egr-1, N = 4 per group. For VMHl egr-1, N = 5 for the Silence/Saline group, and N = 4 for all other groups.

### Statistics

Data processing was conducted in both Python (version 3.7; using Spyder (version 4.0.1; (103)), and the pandas (104) and numpy (105) libraries), and R (version 4.2.0; using R Studio (106) and the tidyverse package (107); all statistical analyses were conducted in R.

To test whether GABA receptor agonist infusion into Field L disrupted the response of NCM to auditory playback, we used a linear mixed model with stimulus firing rate (Hz) as the dependent variable, and recording time course as an independent variable, with auditory stimulus ID as a random effect. Posthoc linear contrasts were evaluated using Tukey’s HSD posthoc tests (emmeans package).

To test whether egr-1 labeling or behavior differed between our experimental treatments, we used 2-way ANOVAs (Anova function in car package), including main effects of song stimulus (i.e., song, or no song) and of cannula infusion treatment (i.e., with GABA receptor agonists baclofen & muscimol, or with saline), and their interaction. We evaluated assumptions of parametric statistics for normality with Shapiro’s test (shapiro.test in stats package), and for equal variances with Levene’s test (leveneTest in car package). Egr-1 cell counts did not meet assumptions for VMHm and BSTm, and were thus log transformed; however, in Figure 3, untransformed data are shown. If significant main effects or interactions were identified in omnibus ANOVA tests, we then conducted Tukey’s HSD posthoc tests (PostHocTest in DescTools).

For comparing the proportion of behaviors done during song playback in birds that received either GABA receptor agonists or saline in auditory pallium, we used Wilcoxon Signed Rank tests. For assessing the relationship between behavior and egr-1 labeling in VMHl, we used a 2-way ANCOVA. When the ANCOVA showed that behavior was a significant predictor of egr-1 labeling, we used a simple linear regression as a posthoc test.

## Acknowledgements

We are grateful to Violet Ivan, Nick Ambrosio, and Britt Mardis for assistance with pilot experiments and data collection. We thank Dr. Andrea Silva-Gotay, Sam Scott, and Dr. Heather Richardson for assistance with SCORE imaging. This work was supported by US National Institutes of Health (NIH) F32 DC018508 and NIH K99 DC020588 awarded to J.S., and NIH R01 NS082179 awarded to L.R.H.

## Author contributions

J.A.S. and L.R.H. designed the research; J.A.S. and A.P.L. performed the research; J.A.S. and A.P.L. analyzed the data; J.A.S. and L.R.H. wrote the paper.

## References

1. S. W. Newman, The medial extended amygdala in male reproductive behavior - A node in the mammalian social behavior network. Ann Ny Acad Sci 877, 242–257 (1999).

2. J. L. Goodson, The vertebrate social behavior network: evolutionary themes and variations. Horm Behav 48, 11–22 (2005).

3. G. D. Fernandez-Fewell, M. Meredith, c-fos expression in vomeronasal pathways of mated or pheromone- stimulated male golden hamsters: contributions from vomeronasal sensory input and expression related to mating performance. J. Neurosci. 14, 3643–3654 (1994).

4. A. L. Falkner, P. Dollar, P. Perona, D. J. Anderson, D. Lin, Decoding Ventromedial Hypothalamic Neural Activity during Male Mouse Aggression. J. Neurosci. 34, 5971–5984 (2014).

5. J. F. Bergan, Y. Ben-Shaul, C. Dulac, Sex-specific processing of social cues in the medial amygdala. eLife 3, e02743 (2014).

6. D. L. Maney, C. T. Goode, H. S. Lange, S. E. Sanford, B. L. Solomon, Estradiol modulates neural responses to song in a seasonal songbird. Journal of Comparative Neurology 511, 173–186 (2008).

7. H. E. Schubloom, S. C. Woolley, Variation in social relationships relates to song preferences and EGR1 expression in a female songbird. Developmental Neurobiology 76, 1029–1040 (2016).

8. J. P. Lorberbaum, et al., A potential role for thalamocingulate circuitry in human maternal behavior. Biological Psychiatry 51, 431–445 (2002).

9. P. M. Forlano, et al., Attention and Motivated Response to Simulated Male Advertisement Call Activates Forebrain Dopaminergic and Social Decision-Making Network Nuclei in Female Midshipman Fish. Integrative and Comparative Biology 57, 820–834 (2017).

10. C. L. Petersen, et al., Exposure to Advertisement Calls of Reproductive Competitors Activates Vocal-Acoustic and Catecholaminergic Neurons in the Plainfin Midshipman Fish, Porichthys notatus. PLOS ONE 8, e70474 (2013).

11. M.-F. Cheng, J. P. Peng, P. Johnson, Hypothalamic Neurons Preferentially Respond to Female Nest Coo Stimulation: Demonstration of Direct Acoustic Stimulation of Luteinizing Hormone Release. J. Neurosci. 18, 5477–5489 (1998).

12. J. S. Lonstein, D. A. Simmons, J. M. Swann, J. M. Stern, Forebrain expression of c-fos due to active maternal behaviour in lactating rats. Neuroscience 82, 267–281 (1997).

13. H. K. Chadha, C. H. Hubscher, Convergence of nociceptive information in the forebrain of female rats: Reproductive organ response variations with stage of estrus. Experimental Neurology 210, 375–387 (2008).

14. H. N. Mallick, S. K. Manchanda, V. M. Kumar, Sensory Modulation of the Medial Preoptic Area Neuronal Activity by Dorsal Penile Nerve Stimulation in Rats. The Journal of Urology 151, 759–762 (1994).

15. U. Mayer, O. Rosa-Salva, G. Vallortigara, First exposure to an alive conspecific activates septal and amygdaloid nuclei in visually-naïve domestic chicks (Gallus gallus). Behavioural Brain Research 317, 71–81 (2017).

16. S. S. Burmeister, E. D. Jarvis, R. D. Fernald, Rapid Behavioral and Genomic Responses to Social Opportunity. PLOS Biology 3, e363 (2005).

17. J. L. Goodson, A. K. Evans, L. Lindberg, C. D. Allen, Neuro–evolutionary patterning of sociality. Proceedings of the Royal Society B: Biological Sciences 272, 227–235 (2005).

18. N. H. Prior, E. J. Bentz, A. G. Ophir, Reciprocal processes of sensory perception and social bonding: an integrated social-sensory framework of social behavior. Genes, Brain and Behavior 21, e12781 (2022).

19. N. E. G. Hoglen, D. S. Manoli, Cupid’s quiver: Integrating sensory cues in rodent mating systems. Front Neural Circuits 16, 944895 (2022).

20. K. S. Lynch, et al., A neural basis for password-based species recognition in an avian brood parasite. Journal of Experimental Biology 220, 2345–2353 (2017).

21. S. J. Chew, D. S. Vicario, F. Nottebohm, A large-capacity memory system that recognizes the calls and songs of individual birds. PNAS 93, 1950–1955 (1996).

22. C. D. Meliza, D. Margoliash, Emergence of Selectivity and Tolerance in the Avian Auditory Cortex. J. Neurosci. 32, 15158–15168 (2012).

23. D. Y. Lin, S.-Z. Zhang, E. Block, L. C. Katz, Encoding social signals in the mouse main olfactory bulb. Nature 434, 470–477 (2005).

24. D. W. Wacker, M. Engelmann, V. A. Tobin, S. L. Meddle, M. Ludwig, Vasopressin and social odor processing in the olfactory bulb and anterior olfactory nucleus. Annals of the New York Academy of Sciences 1220, 106–116 (2011).

25. G. Tasaka, et al., The Temporal Association Cortex Plays a Key Role in Auditory-Driven Maternal Plasticity. Neuron 107, 566–579.e7 (2020).

26. J. K. Schiavo, et al., Innate and plastic mechanisms for maternal behaviour in auditory cortex. Nature 587, 426–431 (2020).

27. B. Cádiz-Moretti, M. Otero-García, F. Martínez-García, E. Lanuza, Afferent projections to the different medial amygdala subdivisions: a retrograde tracing study in the mouse. Brain Struct Funct 221, 1033–1065 (2014).

28. T. Inbar, R. Davis, J. F. Bergan, A sex-specific feedback projection from aromatase-expressing neurons in the medial amygdala to the accessory olfactory bulb. Journal of Comparative Neurology (2021) https://doi.org/10.1002/cne.25236 (October 15, 2021).

29. F. Scalia, S. S. Winans, The differential projections of the olfactory bulb and accessory olfactory bulb in mammals. J. Comp. Neurol. 161, 31–55 (1975).

30. N. Kang, M. J. Baum, J. A. Cherry, A direct main olfactory bulb projection to the ‘vomeronasal’ amygdala in female mice selectively responds to volatile pheromones from males. European Journal of Neuroscience 29, 624–634 (2009).

31. T. Osakada, et al., Sexual rejection via a vomeronasal receptor-triggered limbic circuit. Nat Commun 9, 4463 (2018).

32. X. Zha, et al., VMHvl-Projecting Vglut1+ Neurons in the Posterior Amygdala Gate Territorial Aggression. Cell Reports 31, 107517 (2020).

33. P. Chen, W. Hong, Neural Circuit Mechanisms of Social Behavior. Neuron 98, 16–30 (2018).

34. G. D. Fewell, M. Meredith, Experience facilitates vomeronasal and olfactory influence on Fos expression in medial preoptic area during pheromone exposure or mating in male hamsters. Brain Research 941, 91–106 (2002).

35. J. E. Lischinsky, D. Lin, Neural mechanisms of aggression across species. Nat Neurosci 23, 1317–1328 (2020).

36. K. L. Hoke, M. J. Ryan, W. Wilczynski, Social cues shift functional connectivity in the hypothalamus. Proceedings of the National Academy of Sciences 102, 10712–10717 (2005).

37. I. C. Hall, I. H. Ballagh, D. B. Kelley, The Xenopus Amygdala Mediates Socially Appropriate Vocal Communication Signals. J. Neurosci. 33, 14534–14548 (2013).

38. M.-F. Cheng, J. P. Peng, Reciprocal talk between the auditory thalamus and the hypothalamus: an antidromic study. NeuroReport 8, 653–658 (1997).

39. W. Wilczynski, J. D. Allison, Acoustic Modulation of Neural Activity in the Hypothalamus of the Leopard Frog. BBE 33, 317–324 (1989).

40. J. D. Allison, W. Wilczynski, Thalamic and Midbrain Auditory Projections to the Preoptic Area and Ventral Hypothalamus in the Green Treefrog (Hyla Cinerea). BBE 38, 322–331 (1991).

41. S. Valtcheva, et al., Neural circuitry for maternal oxytocin release induced by infant cries. 2021.03.25.436883 (2021).

42. J. Martin Wild, H. J. Karten, B. J. Frost, Connections of the auditory forebrain in the pigeon (columba livia). Journal of Comparative Neurology 337, 32–62 (1993).

43. G. E. Vates, B. M. Broome, C. V. Mello, F. Nottebohm, Auditory pathways of caudal telencephalon and their relation to the song system of adult male zebra finches (Taenopygia guttata). Journal of Comparative Neurology 366, 613–642 (1996).

44. G. S. Berns, et al., Diffusion tensor imaging of dolphin brains reveals direct auditory pathway to temporal lobe. Proc Biol Sci 282, 20151203 (2015).

45. C. E. Carr, R. A. Code, “The Central Auditory System of Reptiles and Birds” in Comparative Hearing: Birds and Reptiles, Springer Handbook of Auditory Research., R. J. Dooling, R. R. Fay, A. N. Popper, Eds. (Springer New York, 2000), pp. 197–248.

46. J. E. Rose, C. N. Woolsey, The relations of thalamic connections, cellular structure and evocable electrical activity in the auditory region of the cat. Journal of Comparative Neurology 91, 441–466 (1949).

47. B. J. Marlin, M. Mitre, J. A. D’amour, M. V. Chao, R. C. Froemke, Oxytocin enables maternal behaviour by balancing cortical inhibition. Nature 520, 499–504 (2015).

48. K. Krishnan, B. Y. B. Lau, G. Ewall, Z. J. Huang, S. D. Shea, MECP2 regulates cortical plasticity underlying a learned behaviour in adult female mice. Nat Commun 8, 14077 (2017).

49. H. J. Barr, E. M. Wall, S. C. Woolley, Dopamine in the songbird auditory cortex shapes auditory preference. Current Biology (2021) https://doi.org/10.1016/j.cub.2021.08.005 (October 3, 2021).

50. S. MacDougall-Shackleton, S. Hulse, G. Ball, Neural bases of song preferences in female zebra finches (Taeniopygia guttata). Neuroreport 9, 3047–3052 (1998).

51. S. E. Maguire, M. F. Schmidt, D. J. White, Social Brains in Context: Lesions Targeted to the Song Control System in Female Cowbirds Affect Their Social Network. PLOS ONE 8, e63239 (2013).

52. M. Macedo-Lima, L. Remage-Healey, Auditory learning in an operant task with social reinforcement is dependent on neuroestrogen synthesis in the male songbird auditory cortex. Hormones and Behavior 121, 104713 (2020).

53. D. M. Schneider, S. M. N. Woolley, Sparse and Background-Invariant Coding of Vocalizations in Auditory Scenes. Neuron 79, 141–152 (2013).

54. J. M. Moore, S. M. N. Woolley, Emergent tuning for learned vocalizations in auditory cortex. Nat Neurosci 22, 1469–1476 (2019).

55. D. J. Bailey, J. C. Rosebush, J. Wade, The hippocampus and caudomedial neostriatum show selective responsiveness to conspecific song in the female zebra finch. Journal of Neurobiology 52, 43–51 (2002).

56. A. V. Medvedev, J. S. Kanwal, Communication call-evoked gamma-band activity in the auditory cortex of awake bats is modified by complex acoustic features. Brain Research 1188, 76–86 (2008).

57. M. H. Davis, I. S. Johnsrude, Hierarchical Processing in Spoken Language Comprehension. J. Neurosci. 23, 3423–3431 (2003).

58. K. Yu, W. E. Wood, F. E. Theunissen, High-capacity auditory memory for vocal communication in a social songbird. Science Advances 6, eabe0440 (2020).

59. T. Q. Gentner, S. H. Hulse, G. F. Ball, Functional differences in forebrain auditory regions during learned vocal recognition in songbirds. J Comp Physiol A 190, 1001–1010 (2004).

60. S. M. H. Gobes, J. J. Bolhuis, Birdsong Memory: A Neural Dissociation between Song Recognition and Production. Current Biology 17, 789–793 (2007).

61. M. Monbureau, J. M. Barker, G. Leboucher, J. Balthazart, Male song quality modulates c-Fos expression in the auditory forebrain of the female canary. Physiology & Behavior 147, 7–15 (2015).

62. L. Remage-Healey, M. J. Coleman, R. K. Oyama, B. A. Schlinger, Brain estrogens rapidly strengthen auditory encoding and guide song preference in a songbird. PNAS 107, 3852–3857 (2010).

63. K. Sen, F. E. Theunissen, A. J. Doupe, Feature Analysis of Natural Sounds in the Songbird Auditory Forebrain. Journal of Neurophysiology 86, 1445–1458 (2001).

64. Y. Chen, J. T. Sakata, Norepinephrine in the avian auditory cortex enhances developmental song learning. Journal of Neurophysiology 125, 2397–2407 (2021).

65. M. Araki, M. M. Bandi, Y. Yazaki-Sugiyama, Mind the gap: Neural coding of species identity in birdsong prosody. Science 354, 1282–1287 (2016).

66. S. Yanagihara, Y. Yazaki-Sugiyama, Auditory experience-dependent cortical circuit shaping for memory formation in bird song learning. Nat Commun 7, 11946 (2016).

67. Y. Mandelblat-Cerf, L. Las, N. Denisenko, M. S. Fee, A role for descending auditory cortical projections in songbird vocal learning. eLife 3, e02152 (2014).

68. J. M. Wild, The ventromedial hypothalamic nucleus in the zebra finch (Taeniopygia guttata): Afferent and efferent projections in relation to the control of reproductive behavior. Journal of Comparative Neurology 525, 2657–2676 (2017).

69. L. Remage-Healey, N. R. Joshi, Changing Neuroestrogens Within the Auditory Forebrain Rapidly Transform Stimulus Selectivity in a Downstream Sensorimotor Nucleus. J. Neurosci. 32, 8231–8241 (2012).

70. W. Kuenzel, Dual hypothalamic feeding system in a migratory bird, Zonotrichia albicollis. American Journal of Physiology-Legacy Content 223, 1138–1142 (1972).

71. W. J. Kuenzel, Multiple effects of ventromedial hypothalamic lesions in the white-throated sparrow,Zonotrichia albicollis. J. Comp. Physiol. 90, 169–182 (1974).

72. R. M. Hnasko, J. D. Buntin, Functional mapping of neural sites mediating prolactin-induced hyperphagia in doves. Brain Research 623, 257–266 (1993).

73. J. D. Buntin, R. M. Hnasko, P. H. Zuzick, Role of the Ventromedial Hypothalamus in Prolactin-Induced Hyperphagia in Ring Doves. Physiology & Behavior 66, 255–261 (1999).

74. P. Viskaitis, et al., Modulation of SF1 Neuron Activity Coordinately Regulates Both Feeding Behavior and Associated Emotional States. Cell Rep 21, 3559–3572 (2017).

75. B. M. King, The rise, fall, and resurrection of the ventromedial hypothalamus in the regulation of feeding behavior and body weight. Physiology & Behavior 87, 221–244 (2006).

76. W. C. Krause, H. A. Ingraham, “Origins and Functions of the Ventrolateral VMH: A Complex Neuronal Cluster Orchestrating Sex Differences in Metabolism and Behavior” in Sex and Gender Factors Affecting Metabolic Homeostasis, Diabetes and Obesity, Advances in Experimental Medicine and Biology., F. Mauvais-Jarvis, Ed. (Springer International Publishing, 2017), pp. 199–213.

77. M. J. Gibson, M. Cheng, Neural mediation of estrogen-dependent courtship behavior in female ring doves. Journal of Comparative and Physiological Psychology 93, 855–867 (1979).

78. S. A. Heimovics, L. V. Riters, Breeding-context-dependent relationships between song and cFOS labeling within social behavior brain regions in male European starlings (Sturnus vulgaris). Hormones and Behavior 50, 726–735 (2006).

79. S. M. Correa, et al., An Estrogen-Responsive Module in the Ventromedial Hypothalamus Selectively Drives Sex-Specific Activity in Females. Cell Reports 10, 62–74 (2015).

80. A. Petzold, H. E. van den Munkhof, R. Figge-Schlensok, T. Korotkova, Complementary lateral hypothalamic populations resist hunger pressure to balance nutritional and social needs. Cell Metabolism (2023) https://doi.org/10.1016/j.cmet.2023.02.008 (February 27, 2023).

81. Z. Zhang, et al., Estrogen-sensitive medial preoptic area neurons coordinate torpor in mice. Nat Commun 11, 6378 (2020).

82. M. B. Pardi, et al., A thalamocortical top-down circuit for associative memory. Science 370, 844–848 (2020).

83. S. Ono, K. Okanoya, Y. Seki, Hierarchical emergence of sequence sensitivity in the songbird auditory forebrain. J Comp Physiol A 202, 163–183 (2016).

84. C. V. Mello, T. Kaser, A. A. Buckner, M. Wirthlin, P. V. Lovell, Molecular architecture of the zebra finch arcopallium. Journal of Comparative Neurology 527, 2512–2556 (2019).

85. J. R. Merritt, et al., A supergene-linked estrogen receptor drives alternative phenotypes in a polymorphic songbird. Proceedings of the National Academy of Sciences 117, 21673–21680 (2020).

86. M.-F. Cheng, M. Chaiken, M. Zuo, H. Miller, Nucleus Taenia of the Amygdala of Birds: Anatomical and Functional Studies in Ring Doves (*Streptopelia risoria*) and European Starlings (*Sturnus vulgaris*). Brain Behav Evol 53, 243–270 (1999).

87. T. Raam, W. Hong, Organization of neural circuits underlying social behavior: A consideration of the medial amygdala. Current Opinion in Neurobiology 68, 124–136 (2021).

88. R. B. Simerly, L. W. Swanson, The organization of neural inputs to the medial preoptic nucleus of the rat. The Journal of comparative neurology 246, 312–42 (1986).

89. D. Schaeuble, B. Myers, Cortical–Hypothalamic Integration of Autonomic and Endocrine Stress Responses. Frontiers in Physiology 13 (2022).

90. L. A. O’Connell, H. A. Hofmann, The Vertebrate mesolimbic reward system and social behavior network: A comparative synthesis. Journal of Comparative Neurology 519, 3599–3639 (2011).

91. A. M. Kelly, A consideration of brain networks modulating social behavior. Hormones and Behavior 141, 105138 (2022).

92. L. V. Riters, S. J. Alger, Neuroanatomical evidence for indirect connections between the medial preoptic nucleus and the song control system: possible neural substrates for sexually motivated song. Cell Tissue Res 316, 35–44 (2004).

93. L. V. Riters, G. F. Ball, Lesions to the Medial Preoptic Area Affect Singing in the Male European Starling (Sturnus vulgaris). Hormones and Behavior 36, 276–286 (1999).

94. J. A. Boulant, Hypothalamic mechanisms in thermoregulation. Fed Proc 40, 2843–2850 (1981).

95. S. Yu, M. François, C. Huesing, H. Münzberg, The Hypothalamic Preoptic Area and Body Weight Control. NEN 106, 187–194 (2018).

96. J. A. Spool, et al., Genetically identified neurons in avian auditory pallium mirror core principles of their mammalian counterparts. Current Biology 31, 2831–2843.e6 (2021).

97. M. J. Coleman, A. Roy, J. M. Wild, R. Mooney, Thalamic Gating of Auditory Responses in Telencephalic Song Control Nuclei. J. Neurosci. 27, 10024–10036 (2007).

98. C. V. Mello, S. Ribeiro, ZENK protein regulation by song in the brain of songbirds. Journal of Comparative Neurology 393, 426–438 (1998).

99. H. J. Karten, et al., Digital atlas of the zebra finch (Taeniopygia guttata) brain: A high-resolution photo atlas. Journal of Comparative Neurology 521, 3702–3715 (2013).

100. B. Nixdorf-Bergweiler, H. Bischof, A Stereotaxic Atlas Of The Brain Of The Zebra Finch, Taeniopygia Guttata: With Special Emphasis On Telencephalic Visual And Song System Nuclei in Transverse and Sagittal Sections. (National Center for Biotechnology Information, 2007).

101. D. G. Gonsalvez, et al., Imaging and Quantification of Myelin Integrity After Injury With Spectral Confocal Reflectance Microscopy. Frontiers in Molecular Neuroscience 12 (2019).

102. A. J. Schain, R. A. Hill, J. Grutzendler, Label-free in vivo imaging of myelinated axons in health and disease with spectral confocal reflectance microscopy. Nat Med 20, 443–449 (2014).

103. P. Raybaut, Spyder-documentation (2009).

104. W. Mckinney, pandas: a Foundational Python Library for Data Analysis and Statistics. Python High Performance Science Computer (2011).

105. C. R. Harris, et al., Array programming with NumPy. Nature 585, 357–362 (2020).

106. RStudio Team, RStudio: Integrated Development for R (2022).

107. H. Wickham, et al., Welcome to the Tidyverse. Journal of Open Source Software 4, 1686 (2019).

