## Supplemental Figure 1 for "Top-down, auditory pallial regulation of the social behavior network"

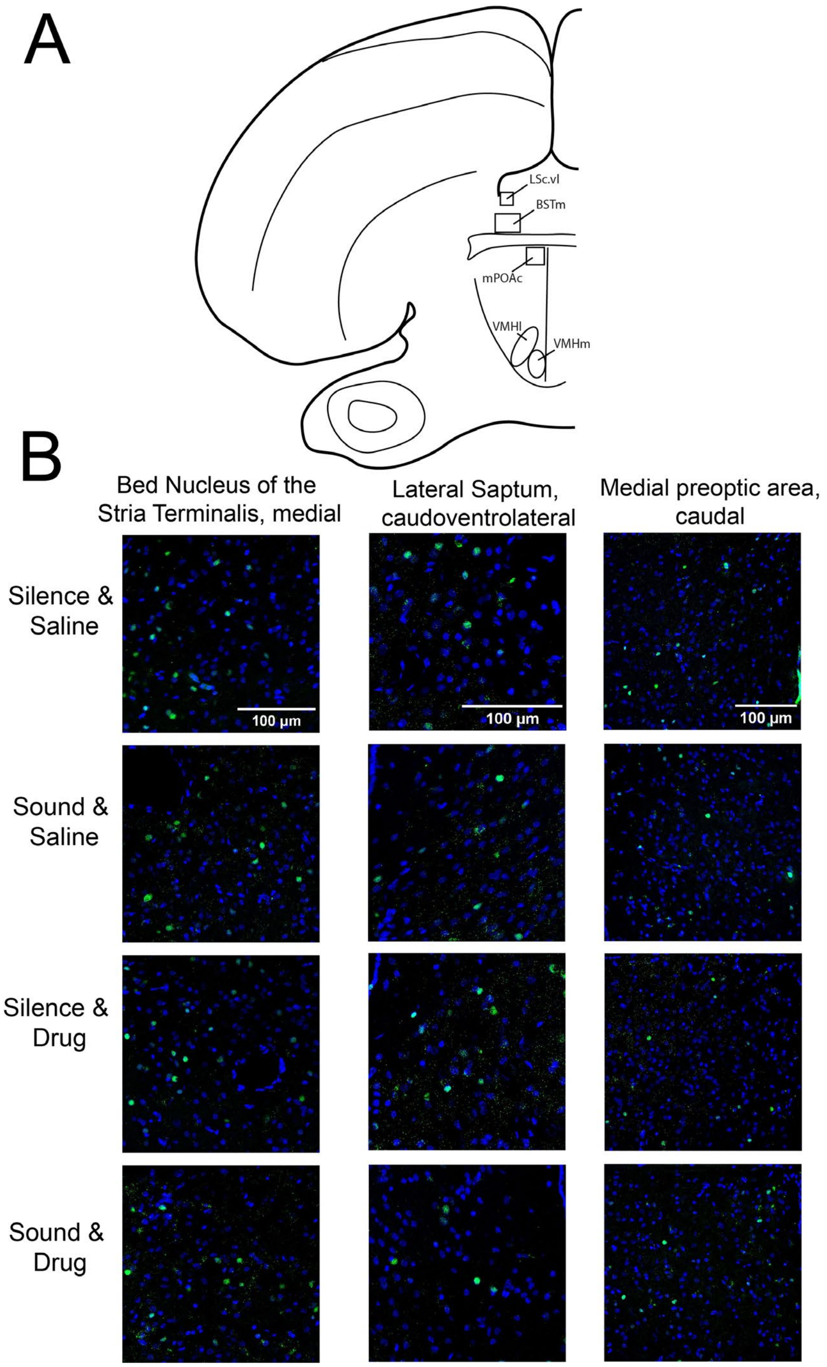


**Figure S1**

(**A**) A hemi-section in the coronal plane at the level of anterior commissure showing locations of brain regions imaged. (**B**) Exemplar images of egr-1 expression (green) and DAPI (blue) in the Bed nucleus of the stria terminalis, medial (BSTm; left column), the Lateral Septum, caudoventrolateral (LSc.vl; middle column), and the medial preoptic area, caudal (mPOAc; right column). Each row represents an experimental treatment group, including whether a bird received a saline infusion or GABA receptor agonist infusion into auditory pallium, and whether a bird was exposed to silence or song playback.
